# Variations in Homoeologous Dosage and Epigenomics Mark the Early Evolution of Synthetic *Brassica* Tetraploids

**DOI:** 10.1101/2023.06.27.543697

**Authors:** Kang Zhang, Yinqing Yang, Lei Zhang, Yinan Cui, Jian Wu, Jianli Liang, Xing Li, Lingkui Zhang, Xin Zhang, Yiyue Zhang, Zhongwei Guo, Shumin Chen, Michael Freeling, Xiaowu Wang, Feng Cheng

## Abstract

Polyploidization is important in plant evolution and is becoming increasingly important in crop breeding and material creation. Studies have provided evidence for structural variations and epigenomic repatterning in synthetic polyploidizations, but the relationships between structural and epigenomic variations, as well as their effects on gene expression and phenotypic variations are unknown. Here, we investigated the genome-wide large deletion/duplication regions (DelDups) and genomic methylation dynamics, in the leaf organ as a representative, of progenies from eight generations that derived from the synthetic tetraploidization between *Brassica rapa* and *Brassica oleracea*. We found that half or complete deletion/duplication of fragments ranging in size from 400 kb to 65.85 Mb, with a mean size of 5.70 Mb, occurred frequently from the first generation of selfing and thereafter. The genes located in these DelDups expressed at levels expected for a positive dosage effect, as indicated by the positive association between expression and the copy number of these genes. Plants containing these DelDups also showed distinct phenotypic variations. The whole genome methylation level experienced significant fluctuations in different generations and eventually decreased in the latter generations. Moreover, the DelDups did not show methylation changes from other individuals of the same generation, and the local regions with methylation alterations did not affect gene expression. Our findings provide new insights into the early evolution of polyploid genomes and guide the use of synthetic polyploidizations in breeding.

## INTRODUCTION

Polyploidization is one of the most important driving forces for the evolution of plants, and all plants evolved from ancient polyploids. Many plant clades experienced multiple rounds of polyploidization events (1, 2). Ancient polyploidizations, also called paleo-polyploidizations, were followed by genome rediploidization, i.e., the polyploidized genomes rearranged and returned to new diploids segregating their chromosomes like diploids (3). The repeated cycles of “polyploidization–rediploidization”, accompanied by extensive genome and gene fractionations, might have given birth to a burst of new species. In addition to paleo-polyploidization, neo-polyploidization refers to events that occurred in relatively recent times, so there has not been enough time for diploidization or fractionation. Synthetic polyploids are the ultimate neopolyploid. Neopolyploid genomes keep multiple sets of chromosomes each set similar to what they were before merging (4).

Neo-polyploidization is frequently observed in species from many plant families. Many neo-polyploidized species have been selected and domesticated as crop species (5), including tetraploid rapeseed and peanut, hexaploid wheat and kiwi fruit, and octoploid sugarcane and strawberry (6–11). Neo-polyploid plants can have many advantages, including combined traits of both elite parents, fixed hybrid vigor of distant parents, enhanced defense to biotic and abiotic stress, increased yield, and the creation of novel features or materials that otherwise would not exist (4, 12, 13). Polyploidization—both autopolyploidization (duplication of one genome) and allopolyploidization (merging of different genomes, like what we study in this report)—is being increasingly adopted by breeders and consciously applied in crop breeding. Nevertheless, there are important issues that should be noted in utilizing neo-polyploids in breeding projects, such as genomic shock and aneuploidy occurring in the progeny of synthetic neo-polyploids, which might hinder the use of these materials. These issues associated with neo-polyploidization should be thoroughly investigated to make better use of polyploidization in breeding programs.

Allopolyploidization and hybridization between distant genomes tend to introduce genomic shock into the progeny (14). Genomic shock, while originally ascribed to the results of repeated chromosomal breaks, now refers to any saltatory event that might initiate a series of genome-wide alterations, including DNA methylation changes, reactivation of transposable elements (TEs), genetic instability or chromosomal rearrangements (14). As previously observed, allopolyploidization is often accompanied by genome-wide alterations in DNA methylation. The level of DNA methylation was found to be higher in natural (e.g., tobacco (15)) and resynthesized allopolyploids (e.g., *Arabidopsis* allotetraploids (16)) than in their parents. It has also been reported that allopolyploidization can trigger the additive expression of methyltransferase genes, thus maintaining a high methylation levels of polyploidized genomes (17). High levels of methylation, especially of the TE sequences, are important in “harmonizing” the genome structure and gene expression (18). The reactivation of TEs is also associated with genomic shock. These activated TEs have important effects on genome reorganization and gene functional changes. TE sequences could be reactivated in the offspring of interspecific hybrids and allopolyploids (19). They are randomly inserted into the host genome and might participate in the subsequent regulation of gene expression (20). Meanwhile, the reactivation of TEs might play an important role in promoting genome rearrangement through genomic transposition or recombination (21). Chromosome rearrangements are another major manifestation of genomic shock. The content of rearrangements by hybridization and allopolyploidization varies substantially across species. For example, massive genomic rearrangements have occurred in *Tragopogon* and *Brassica* allopolyploids (22, 23), whereas the genome of synthetic cotton has retained the structure of parental chromosomes (24).

Aneuploidy and the following genetic instability of individual chromosomes or genomic fragments, which may be associated with the chromosomal consequences of genomic shock, are severe problems of neo-polyploidization, especially for the breeder. Neo-polyploids often appear to be accompanied by chromosome number changes (25) due to inter-genomic parental conflicts, especially in allopolyploids. The changes that can occur on the level of complete chromosomes or chromosomal fragments, with much variation among species (26). For example, wheat allotetraploids display higher levels of aneuploidy (27) than the others (24). These wheat aneuploidy events generally occurred in the first generation of hybrids of allotetraploids (27), compared to aneuploidy in other allotetraploids, as assayed using chromosomal molecular marker or karyotype analysis (28, 29). Additionally, early aneuploids showed larger variations in homologous chromosomes than in nonhomologous chromosomes (28, 29). Homoeologous exchange is one of the main ways to generate aneuploids, through which unusual synapsis and recombination of homoeologous chromosomes in allotetraploids occurred during meiosis, which then produces sperm or egg cells with unusual haplotypes for aneuploids. There are genomic fragments that showed copy number variations in Aneuploids. The general rule for a fragment dosage change is expression dosage effect for genes on that fragment, such as what had been observed in maize (30). However, the gene balance hypothesis raised by Birchler *et al*. (31, 32) indicates that genes on chromosomal segments can show dosage, compensation or inverse dosage effects, which obscured the general expectation of dosage effect. Moreover, it is important to note that aneuploidy contributes to phenotypic variations and the rapid morphological divergence of plants in the progeny population derived from new polyploidization (33). Studies have suggested that aneuploidy has a significant impact on phenotypic variability, including sterility (34), flowering time (35), and disease resistance (36), by introducing alterations in gene expression via gene dosage effects/compensation and by break-associated position effects (28).

In addition to the phenotypic nonuniformity introduced by genomic shock and aneuploidy in synthetic polyploid populations, phenotypic instability has also been observed in breeding studies using new polyploids. Phenotypic instability can be manifested as variable phenotypes among different generations and incomplete penetrance of the traits of interest. For example, some traits are always lost or not consistently inherited by progeny during self-crossing. The variations in phenotypes or inheritance of the target traits usually vary from the classical Mendelian genetic model. Nonuniformity and instability of phenotypes may sometimes hinder the application of polyploids in breeding programs. Therefore, further investigation is needed to explore the underlying genomic mechanisms. Knowing mechanisms should enhance breeding applications using new polyploids.

In this study, we systematically investigated the variation patterns of genomic fragments, gene expression, and DNA methylation in each of eight consecutive inbreeding generations of synthetic *Brassica* allotetraploids originating from the synthetic wide hybrid *Brassica rapa* × *Brassica oleracea*, as well as in specific cultivars of natural *B. napus*. We studied the leaf organ as representative of the plant. We paid special attention to individual chromosome and genomic fragments that showed aneuploidy to explore the potential mechanisms associated with phenotype nonuniformity and instability of the progeny of new polyploids. We found that different levels of copy number variations in homoeologous fragments frequently occurred in the self-crossed progeny of the synthetic polyploid. The expression of genes—assayed in leaf organ—located in these fragments showed a dosage effect. Because these fragments containing “extra” copies of genes also displayed extra levels of mRNA, such dosage effects might be related to the mechanism underlying associated changes in phenotype. We also found whole genome methylation fluctuations among different generations, although these large-scale DNA methylation changes did not have a detectable impact on gene expression. Moreover, the homoeologous fragments with copy number variations did not show clear alterations in sequence methylation. Our study of the whole genome structure and methylation variations in synthetic polyploids provides valuable results that could guide the utilization of synthetic polyploidization in crop breeding.

## RESULTS

### Homoeologous Copy Number Variations Occurred after Self-crossing in Synthetic Tetraploids

Remarkable large deletion/duplication regions (DelDups) were observed in the tetraploid progeny of synthetic *B. napus*. Synthetic tetraploids emerging from *B. rapa* Chiifu (*rapa*, 2*n* = 20) and *B. oleracea* JZS (*oleracea*, 2*n* = 18) (**Methods**). Briefly, the F_1_ was created from the distant hybrid (haploid) of *rapa* and *oleracea,* and the F_2_ by genome doubling (tetraploid, 2*n* = 4*x* = 38) of F_1_, which was induced by colchicine treatment. F_3_ to F_8_ were generated through continuous self-crossing, i.e. these are selfing generations (S_1_ to S_6_). To investigate genomic variations during the process of synthetic tetraploidization, we mapped the resequencing data of individuals from F_1_ to F_8_ onto the merged genome references of *rapa* (Chiifu) and *oleracea* (JZS). We used a sliding window (300 kb window with 30 kb increment) method to calculate the average depth of mapped reads in the local genomic regions of each sample (**Methods**). We found five peaks in the frequency distribution of read mapping depth in self-crossed generations F_3_ to F_8_, in contrast to only one peak in the genomes of the two parents, as well as the F_1_ and F_2_, which had not experienced self-crossing. Using sample F_6_A (A–C represent the three individuals in a generation) as an example (**Figure 1a**), the first peak represents the lowest read depth—almost no reads mapped—indicating completely deleted genomic fragments; the second peak represents the lower read depth—these sliding windows may be located at the genomic regions with half of them deleted, that is, one copy of a pair of homologous chromosomal fragments was lost; the third peak corresponds to a normal level of read depth (similar with the peak in parents, F_1_ and F_2_); the fourth and fifth peaks represent higher and the highest read depth, which likely correspond to the half and fully duplicated genomic regions, respectively. These peaks represent five types of copy number of fragments in tetraploidized genomes (**Figure 1a** and **Supplementary Figure 1**).

**Figure 1.**
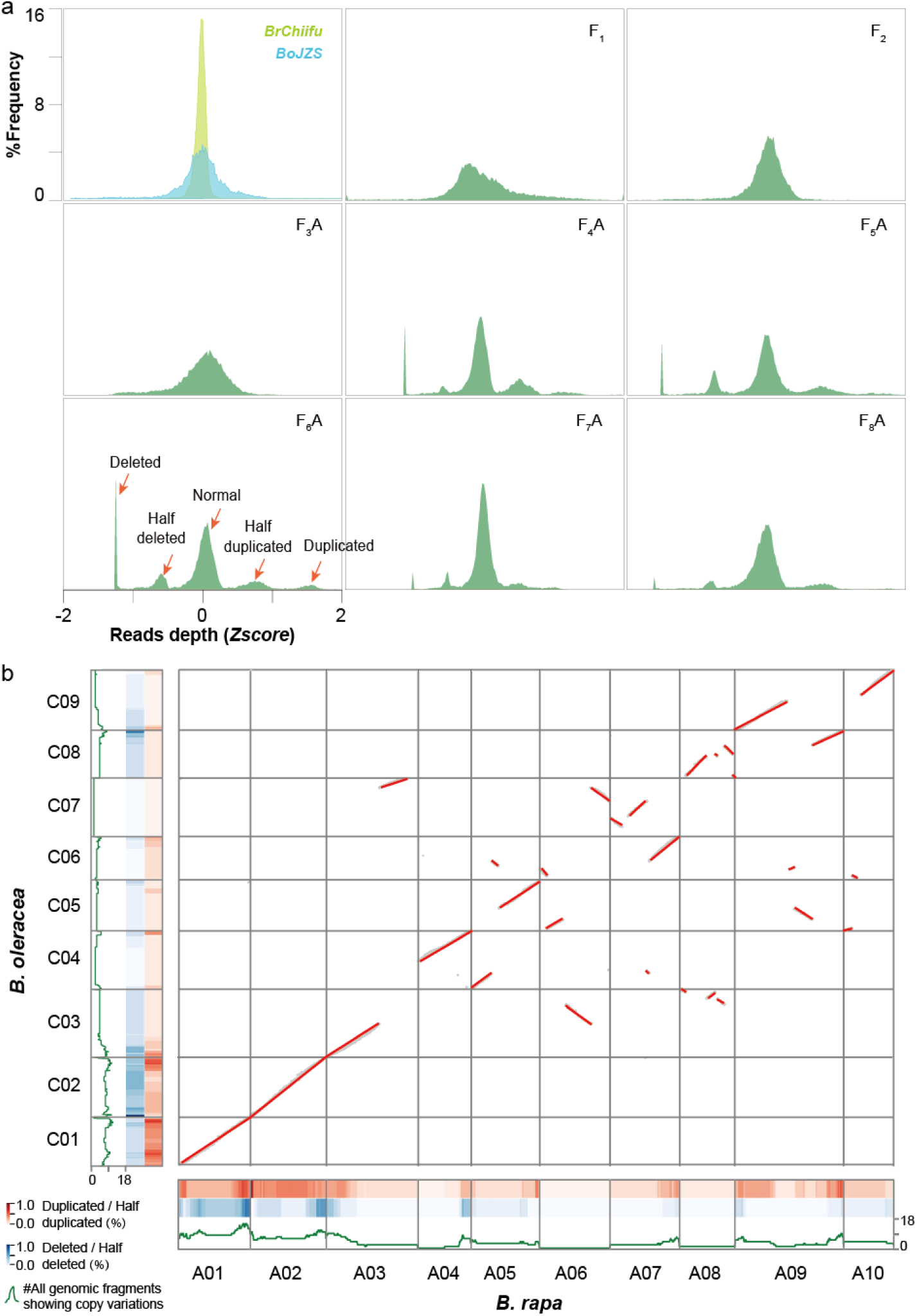
The variations of reads depth and their distribution across the two parental genomes. (**a**) The frequency distribution of read depth in a 200 kb window sliding across the genomes of two parents and from F_1_A to F_8_A. (**b**) The density of genomic regions that showed the reads depth variations in all samples analyzed, which are correlated with genomic synteny.

We separated the genomic regions into four types, which corresponded to the four peaks of read depth (except the normal peak). We obtained 47, 61, 77, and 47 genomic regions (232 in total) representing the deleted, half deleted, half duplicated, and duplicated fragments, respectively, in all of the analyzed samples, with 11.60 (77.45 Mb) such DelDups, on average in each sample, respectively (**Supplementary Table 1**). Among these tetraploid progenies, F_6_A obtained the most (23) DelDups, while F_3_A and F_3_B had no DelDup. We further found many more DelDups (135, accounting for 58.19% of all of the DelDups) located on chromosomes A01, A02, and the head of A03 in *rapa*, as well as C01, C02, and the head of C03 in *oleracea* (**Figures 1** and **2**) than other chromosomes, while no DelDups were found in A06 and C07. In general all fragments of *rapa* and *oleraceae* show both exon and total sequence colinearity (synteny), but some chromosomes have suffered inversions and or translocations. These double chromosomal break aberrations we easily visualize (Fig. 1b) in a nucleotide-nucleotide, all chromosome, nucleotide sequence dot-plot. In such a plot, chromosomal inversions are recognized as short lines at right angles to the no-breakpoint control. Translocations are recognized by finding a fragment of chromosome unexpectedly attached to a new centromere (E.g., Fig. 1b, *rapa* A03 is split over *oleraceae* C03 and C07). Intriguingly, chromosomes with higher DelDup frequencies showed better genomic synteny between *rapa* and *oleracea* than the other chromosomes (**Figure 1b)**. For example, the two longest chromosomes have no breakpoints and have the most fragment copy number variation. However mysterious, the mechanism must be to explain how the most perfectly aligning the chromosomes the higher the number of DelDup, our result is consistent with data from previous reports (28, 29). Specifically, from our results, there were 119 DelDups in *rapa*, with 18 and 28 deleted and half deleted, while 45 and 28 were half duplicated and duplicated, respectively. Meanwhile, there were 113 DelDups in *oleracea*, with 29 and 33 being deleted and half deleted, respectively, while 32 and 19 were half duplicated and duplicated, respectively. The overall comparative number of low (108 in total) and high (104 in total) copies indicated that these DelDups were generated by homoeologous exchanges (HEs) (37) between syntenic fragments of the *rapa* and *oleracea* subgenomes. However, there were more half duplicated or duplicated fragments were found in *rapa* (73) than in *oleracea* (51), while more half deleted or deleted fragments were found in *oleracea* (62) than in *rapa* (46), suggesting to the authors that homoeologous exchanges might be biased toward *rapa*. However, this is inconsistent among different chromosomes. For example, *rapa* had more fractionated fragments in A01 (14) than that of its syntenic chromosome C01 (7) in *oleracea*.

**Figure 2.**
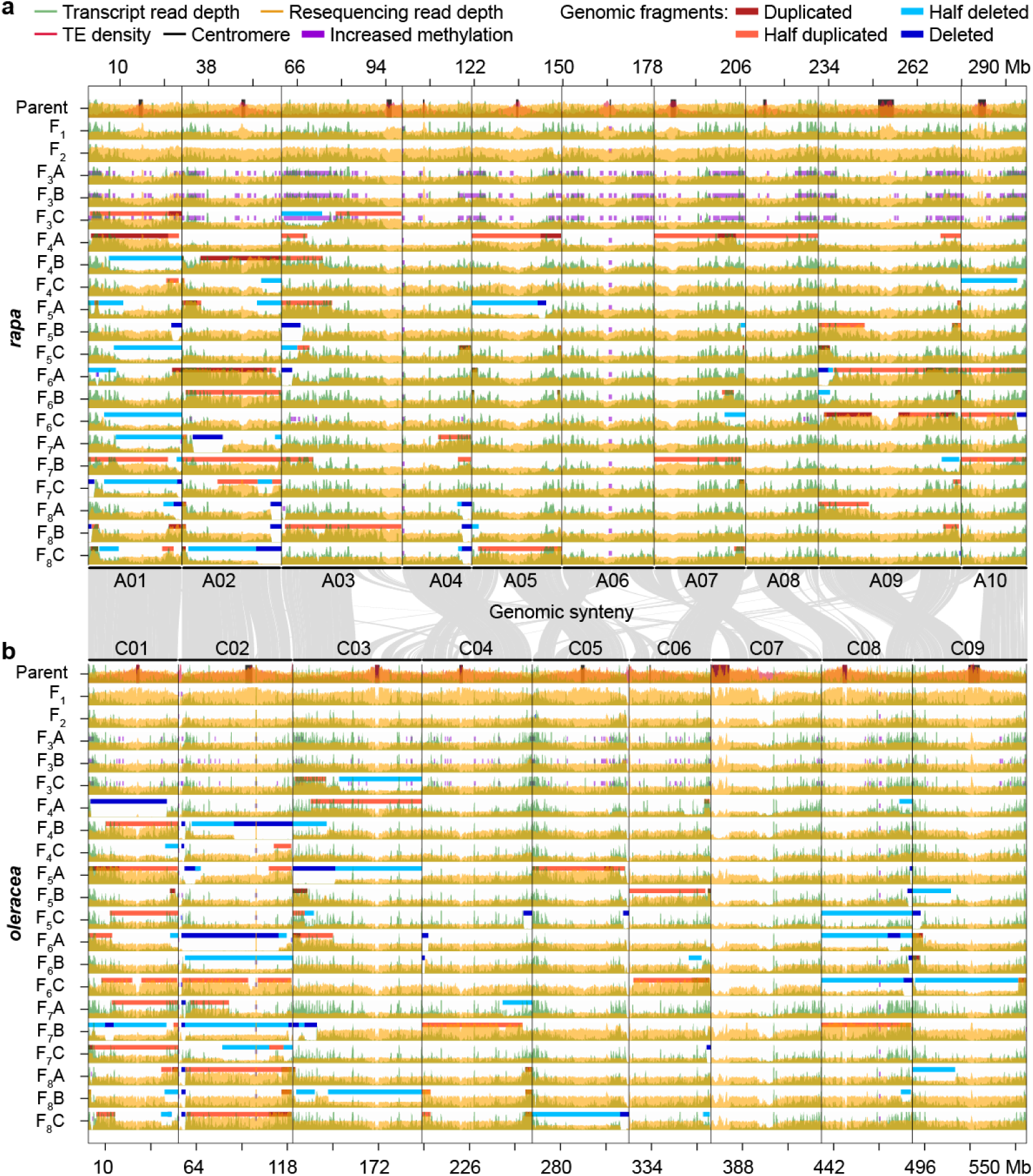
The full landscape of the methylation increased/decreased genomic regions in the hybrid/tetraploid and their self-crossing progenies comparing to their parents, and the distribution of genome-wide transcription quantities. (**a**) and (**b**) denote the comparisons and statistics in the genome/subgenome *rapa*, and the genome/subgenome *oleracea*, respectively.

### Accumulated Dosage of Gene Expression in the DelDup Regions of Aneuploids

Genes located at the DelDups showed strong expression variations as compared to genes not at the DelDups. To evaluate the pattern of expression variations of DelDup-associated genes, we performed transcriptome analysis of these samples. The reads from RNA-seq were mapped to the two parental genomes, and the read depth was calculated using the window-sliding method (**Methods**). As shown in **Figure 2**, the DelDup regions showed clear differences in mapping read abundance, especially for the deleted fragments, which is similar to that of resequencing read mapping and supports the reliability of the DelDups identified. We further investigated the relationships between DelDups and gene transcription by comparing the expression variations of genes located in each of the four groups of DelDups. The expression fold changes of genes located at DelDups were calculated using a comparison between the individual with this DelDup and the other two individuals without this DelDup in the same generation. As shown in **Figures 3a** and **3b**, the deletion of genomic regions was supported by no expression of genes located at these fragments, while for genes located at the half deleted regions, the expression fold-change was ∼0.5, supporting half deletion of these homologous fragments. Genes located at the half duplicated and duplicated regions showed wider ranges of fold-change expression; their expression generally increased to ∼1.5- and ∼2.0-fold, respectively, supporting half and fully duplicated homologous fragments in corresponding genomic regions. These results suggest a strong positive association between the absolute expression dosage of genes and the copy number variations of fragments introduced by homoeologous exchanges. That is our results are expected with 100% dosage effect.

**Figure 3.**
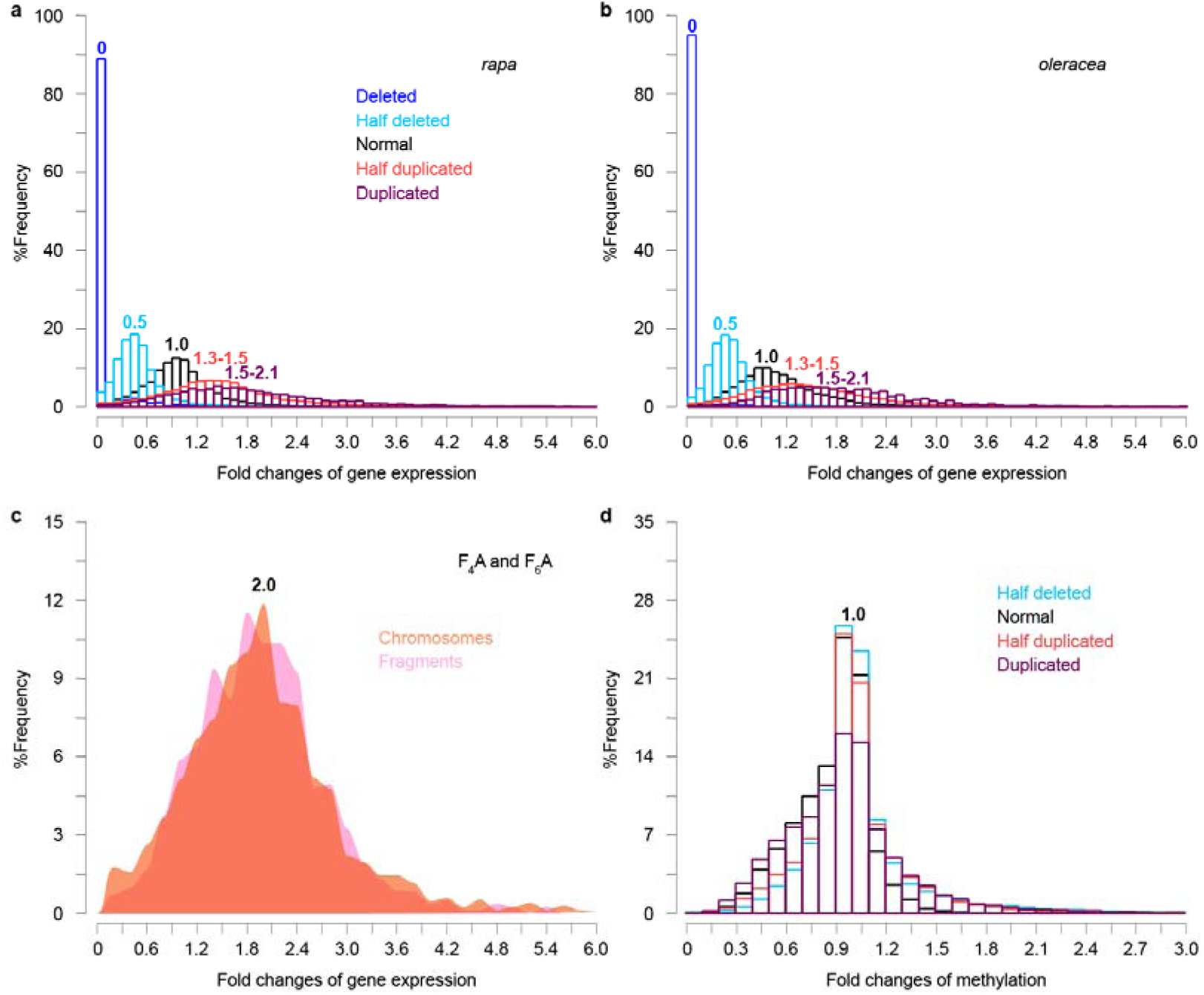
The frequency distribution of fold changes on expression and methylation of genes located at genomic regions with copy number variations. (**a**) The frequency distribution of fold changes of gene expression in subgenome *rapa*. (**b**) The frequency distribution of fold changes of gene expression in subgenome *oleracea*. (**c**) The frequency distribution of fold changes of genes located at the complete chromosomes and local genomic fragments with highest copy abundance. (**d**) The frequency distribution of fold changes of gene methylation in subgenome *rapa*.

Chromosome level and local fragment level DelDups showed similar dosage effects on gene expression. Using samples F_4_A and F_6_A as examples, as shown in **Figure 2**, more than 80% of A01 in F_4_A and A02 in F_6_A were fully duplicated (the highest copy), and both covered the centromere regions. There were two and three local genomic fragments (< 30% of corresponding chromosomes) that were fully duplicated in F_4_A and F_6_A, respectively, both without centromere regions (**Figure 2**). The expression of genes located at the two chromosome-scale DelDups increased by ∼2 times compared to those in the other two individuals from the same generation but without the DelDups. These same results were found for genes located in the five local genomic regions (**Figure 3c** and **Supplementary Figure 2a**). These findings indicate that the duplications of both the chromosome region with centromeres and the genomic fragments without centromeres had the same near-perfect dosage effect on gene expression. Considering that genomic fragments without centromeres are likely lost during mitosis, these duplicated local fragments were likely integrated into homoeologous chromosomes with centromeres through homoeologous exchanges.

Consistent with the DelDups and their impacts on gene expression, we observed phenotypic variations in the corresponding individual plants from F_4_ to F_8_. To determine whether the tetraploid progeny whole plants were phenotypically biased toward the parent *rapa* or *oleracea*, we roughly evaluated four leaf traits known to be different between *rapa* and *oleracea*: 1) leaf trichome (knotted or not), 2) leaf wrinkle (rugose or not), 3) leaf color (dark or light), and 4) leaf serration (serrated or not). We observed that phenotypic bias did exist, and more interestingly, the bias varied in individuals from within the same generation and also among different generations (**Supplementary Figure 3**). For example, in F_4_ there was one individual plant judging from leaves, biased toward *rapa*, one biased toward *oleracea*, and the third one showing a mixed phenotype of both parents; in F_7_, one individual was biased toward *rapa* and two were biased toward *oleracea*. These phenotypic variations might be explained by the large-scale gene expression dosage variations caused by DelDups in synthetic tetraploid progeny generated during self-crossing.

### Genome-wide Methylation Fluctuations in Synthetic Tetraploids

To investigate the patterns of methylation variations during the process of allotetraploidization and subsequent inbreeding. We performed whole genome bisulfite sequencing on the two parents and their progeny from F_1_ to F_8_. Three individual plants were assayed for each generation. Reads of bisulfite sequencing were mapped to the reference genomes of *rapa* and *oleracea* (38, 39) to determine the methylation status of the three contexts of cytosine (C) bases: CpG, CHG, and CHH (40). Subsequently, a sliding window method was used to calculate the weighted methylation level in each genome (**Methods**). We observed whole-genome methylation fluctuations in the original hybrid, and after tetraploidization, and in all generations of the subsequent self-crossing processes (**Figure 4a** and **Supplementary Table 2**). Compared to the two parents, the overall methylation level increased in F_1_, especially in the *rapa* subgenome. In addition, the highly methylated regions surrounding the centromeres were expanded greatly (**Supplementary Figure 4**). However, the methylation level was then decreased in F_2_ in both the *rapa* and *oleracea* subgenomes to an even lower level than that of the two parental genomes. This decrease was not permanent. Interestingly, the high methylation pattern recovered in F_3_, to an even higher level than that of F_1_, especially in the *rapa* subgenome (**Figures 4b** and **4c**, and **Supplementary Figure 4**). This was exemplified by the highly methylated regions surrounding the centromeres, which were larger than those observed in F_1_. The F_3_ plants attaining the highest methylation level were in the first generation of self-crossing. Subsequently, in generations from F_4_ to F_8_, the methylation level was generally decreased compared to that in F_3_ (**Supplementary Figures 5–10**).

**Figure 4.**
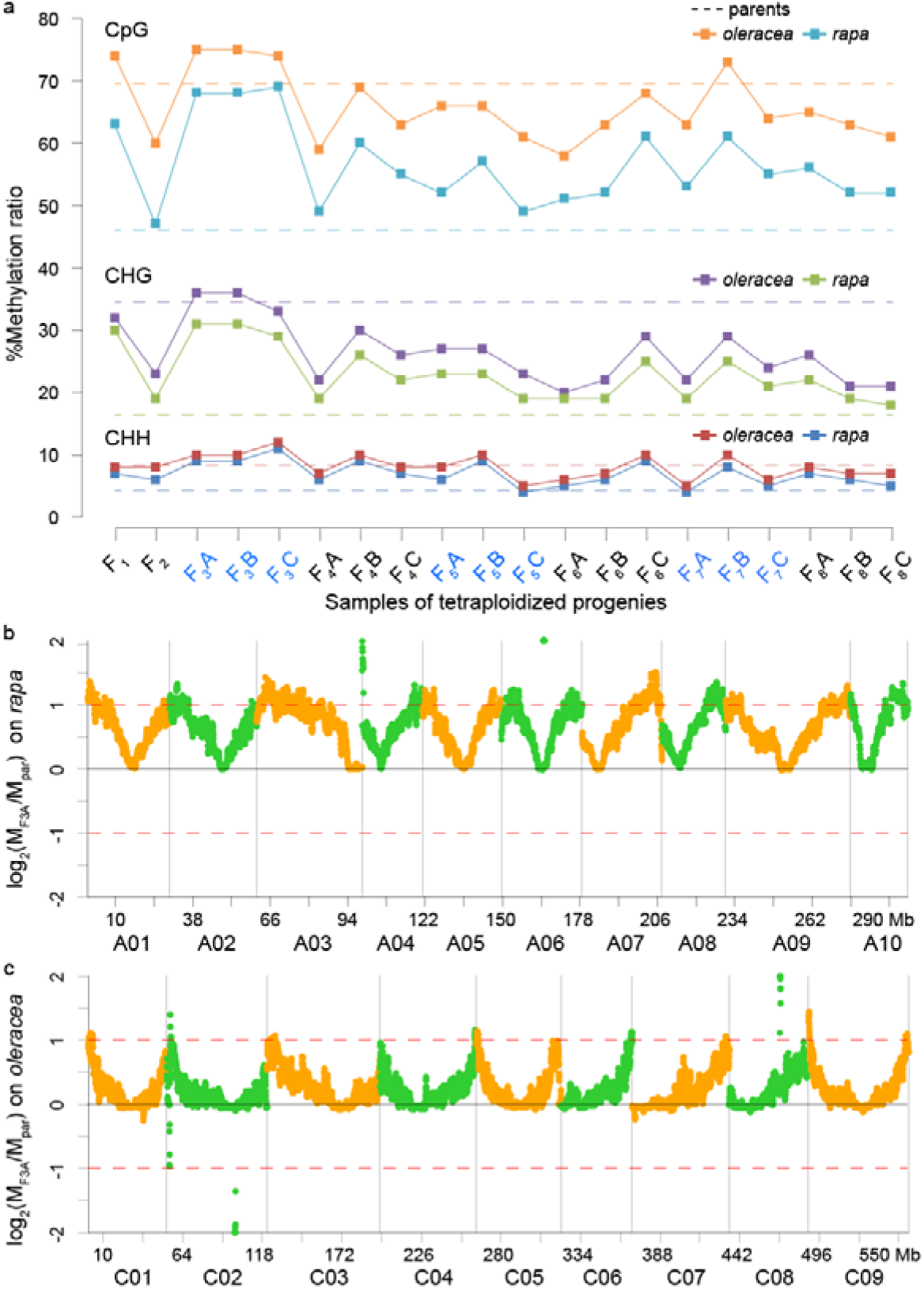
The whole-genome methylation fluctuations among parents and progenies after the merging of two distant parental genomes. (**a**) The comparisons of the whole-genome methylation levels among the F_1_ to F_8_ progenies, as well as the parents (dashed lines), using all the three groups of cysteine locus. (**b**) and (**c**) show examples of the fold changes of genome-wide methylation levels in F_3_A against the two parents *rapa* and *oleracea*. The methylation levels in each of the two parents and the two parental subgenomes in F_3_A were calculated, and then compared using a sliding window of 500 kb that walk across the whole genome. The red dashed-lines denote the threshold of two-fold changes.

The leaf genome-wide methylation alteration did not have any clear impact on the RNA levels transcribed from the synthetic tetroploid genome or any of the inbred generations (**Figure 2**). We performed genome-wide screening to identify local genomic regions with increased or decreased methylation in samples from F_1_ to F_8_. A window-sliding method was used with two-fold changes in methylation levels among F_1_ to F_8_ samples and the parents as a cutoff (**Methods**). In total, we obtained 349 increased methylation regions, with 189 and 160 regions in the two subgenomes, *rapa,* and *oleracea*, respectively (**Figure 2** and **Supplementary Table 3**); most of these (313, 89.68%) were large and continuous genomic regions showing increased methylation in samples of F_3_, the first generation obtained by self-crossing, especially in subgenome *rapa* (**Figure 2**). This was consistent with the increased whole-genome level of methylation in F_3_ as previously presented (**Figures 4b** and **4c**). There were 17 decreased methylation regions that were identified only in *oleracea*. We analyzed gene expression (RNA-seq) in the genomic regions that showed increased or decreased methylation in samples from F_4_ to F_8_ and found no evidence indicating either weakened or enhanced gene expression (**Supplementary Table 4**).

Interestingly, in addition to regions only detected in F_3_, we found four increased methylation loci (corresponding to the 36 increased methylation regions) and one decreased methylation locus (corresponding to the 15 methylation decreased regions) that were shared by multiple tetraploid genomes (≥ 4) (**Supplementary Table 5**). This would be explained if some fraction of methylation pattern were heritable one inbreeding generation to the next. One of the four increased methylation regions was located at the centromere region of chromosome A06. All four of the increased methylation regions occurred in F_1_ or F_2_, while the one decreased methylation region was observed first in F_3_. We further investigated the TE composition in the five changed methylation regions and found that Gypsy long terminal repeat (LTR/Gypsy) was highly enriched in the decreased methylation region, the ratio of which (65.58%) was much higher than the genome background (24.07%) (**Supplementary Table 5**). Meanwhile, there was less LTR/Gypsy (2.18%, 20.53%, and 20.57%) for the three regions with increased methylation, except for the one located at the centromere region (51.68%). These results indicate that, perhaps the decreased methylation region may be associated with activation of LTR/Gypsy elements after tetraploidization in synthetic tetraploid and subsequently inbred genomes.

### Decrease in Whole Genome Methylation Level after Tetraploidization

The methylation level was higher (9.50% in average) in subgenome *oleracea* than in *rapa*, with the methylation level of subgenome *rapa* being higher (9.52% in average) than the parent, while the methylation level of subgenome *oleracea* was generally lower (3.69% in average) than the parent, except for the first self-crossing generation, F_3_ (**Figure 4a** and **Supplementary Table 2**). The oilseed crop *B. napus* was naturally tetraploidized by merging *B. rapa* and *B. oleracea* tens of thousands of years ago. As an annual plant, *B. napus* has experienced tens of thousands of generations to date (41). We selected the *B. napus* cultivar ‘Zhongshuang 11’ (ZS11) as a material to investigate methylation features after long-term evolution (42, 43). Within the ZS11 genome there was also a higher methylation level in subgenome *oleracea* than in subgenome *rapa* (**Supplementary Table 2**); we used the whole genome bisulfite sequencing data for ZS11, data acquired using very similar methods as used for our own synthetic tetraploids. More importantly, as listed in **Supplementary Table 2**, the *rapa* and *oleracea* subgenomes in *B. napus* ZS11 had lower methylation levels than subgenomes *rapa* and *oleracea*, respectively, in our synthetic tetraploid progeny. We evaluated the methylation levels of the TEs. The methylation levels around and in TEs were generally decreased from F_4_ to F_8_, while TE methylation was even further decreased in the genome of ZS11 (**Figure 5**). Taken together, these findings support the conclusion that the genome-wide methylation level decreased for both subgenomes through continuous self-crossing following the tetraploidization event, while the methylation difference between subgenomes *rapa* and *oleracea* remained through tens of thousands of generations to current cultivars of *B. napus*.

**Figure 5.**
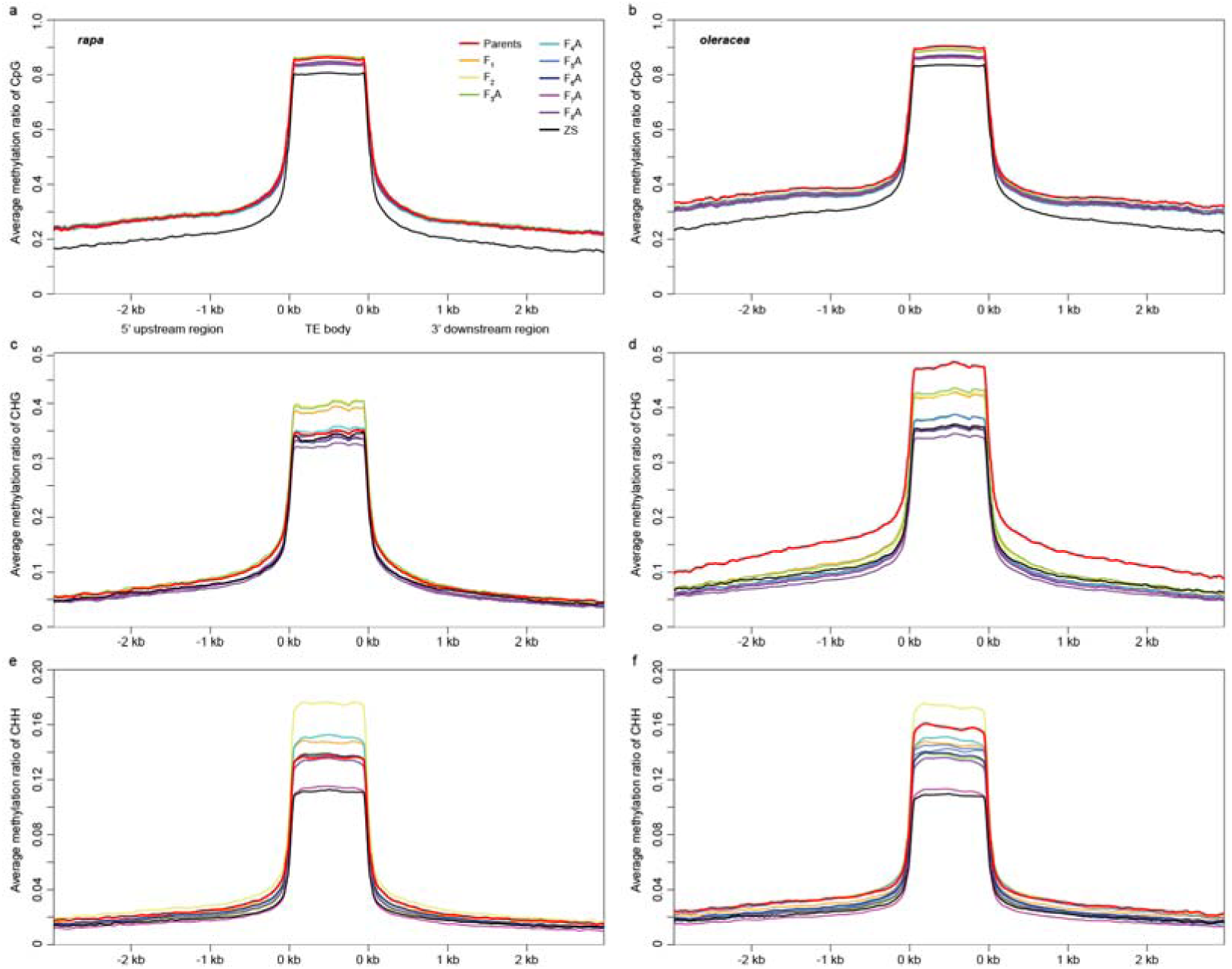
The methylation pattern in the TE bodies and their flanking regions. The methylation ratio in average in and around the TEs of the two parents and the merged progenies F_1_ to F_8_, as well as the *B. napus* cultivar “Zhongshuang 11” in the CpG loci of the parental genome *B. rapa* and sub-genome *rapa* (**a**) and the parental genome *B. oleracea* and sub-genome *oleracea* (**b**), in the CHG loci of the parental genome/sub-genomes *rapa* (**c**) and *oleracea* (**d**), as well as in the CHH loci of the parental genome/sub-genomes *rapa* (**e**) and *oleracea* (**f**).

## DISCUSSION

Synthetic polyploidization has been used as an efficient strategy for generating new crop diversity. To understand the mechanisms underlying this strategy in breeding programs, it is important to consider how the new polyploid genomes behave in terms of genetics and epigenetics. Accordingly, we investigated genome-wide DelDups and DNA methylation densities in the original synthetic tetraploid and in the following inbreeding generations. The original wide hybrid was between representative varieties of two different species *Brassica rapa* (Chinese cabbage) and *Brassica oleraceae* (head cabbage). We revealed the associations between DelDups, gene expression, DNA methylation in all contexts overall, near centromeres, and on TEs, and phenotypic variations over the consecutive inbreeding generations. These findings contribute to our understanding of genome evolution in early polyploidization and could guide the utilization of synthetic polyploidization in breeding applications.

### Extensive Homoeologous Exchanges Altered the Genetic Composition of Polyploids

The doubling of two distant hybrid genomes resulted in two sets of homologous chromosomes co-existing in one genome, i.e., allotetraploidization. The new genome, with its two subgenomes—is stable because it is a double haploid (amphidiploid). Theoretically, both parental genomes, now subgenomes, are homozygous in this new tetraploid genome. It means that the new tetraploid is genetically fixed, and will not “segregate away” through self-crossing. Therefore, plants of the progeny population from tetraploidization are expected to be phenotypically identical. That expectation was not met. Clear phenotypic variations were observed in individuals generated from the self-crossing of our synthetic polyploids, with some phenotypically biased toward parent *B. rapa* and some biased toward parent *B. oleracea*. Considering that these individual plants were grown in the same environment, the phenotypic differences should reflect genomic variations, like mutations or chromosomal aberrations.

Through mapping of resequencing reads, we observed remarkable variations in read depth in large genomic regions that occurred in generations F_3_–F_8_ of the synthetic tetraploids compared to those of the parents. These large genomic regions contain copy number variations of the corresponding fragments. Considering that the homoeologous compensation relationships of deletion and duplication between subgenomes *rapa* and *oleraceae*, these DelDups were deduced to be formed by homoeologous exchanges. Transcription of genes located at these DelDups showed a dosage effect: the more copies of a fragment caused higher expression (summed expression of multiple gene copies) of the genes in the fragment. Additionally, no methylation variations were observed in these DelDup regions, indicating that typical epigenetic silencing was not involved in homoeologous exchanges events.

Genomic variations that occurred from the first self-crossed generation (F_3_) likely play an important role in promoting phenotypic variations of synthetic polyploids, although it is hard to trace the causal genes of these phenotypic differences due to the large-scale changes in homoeologous exchanges. Taken together, the DelDups and resultant gene expression dosage variations are candidate drivers of phenotypic diversity in the progeny population of new polyploids (28).

These DelDups might contribute, like other sorts of mutation, to phenotypic plasticity and innovation, which are then subjected to natural selection during polyploid evolution. Considering that ancient polyploids largely survived drastic environmental changes or natural disasters, the DelDups in polyploids and the resulting genetic diversity may play an important role in adapting to rapidly changing environments and are thus beneficial to the successful survival of plants.

These findings raise an issue that should be considered in breeding programs using synthetic polyploids. DelDups showed an obvious positive correlation with phenotypic diversity in synthetic polyploid populations. Breeders can work with all sorts of variation. However, when the breeding aim is to fix hybrid vigor in synthetic polyploids, these DelDups may introduce unexpected variables, such as differences in the genetic identity and performance of the polyploid population. To obtain a synthetic polyploid population without DelDups, multiple generations of out-crossing are therefore suggested to remove DelDups, which may mimic the situation of wild polyploid populations.

### Large-scale DNA Methylation Level Changes among Synthetic Tetraploid Plants and Clades did not Noticeably Impact Gene Expression

Through the combination analysis of genomic methylation and gene expression (via RNA-seq in leaves) between tetraploid progeny and their parents, we revealed that the expression of genes located at the 349 regions showing increased methylation was no different than regions in the parents or in any other region. Based on the aforementioned results, these 349 regions can be grouped into two types: 1) samples with a genome-wide methylation increase, which was true for the 313 increased methylation regions in the genomes of the three individual plants from generation F_3_; and 2) genomic regions, whose methylation increases are shared in multiple samples. We found four such shared regions in chromosomes A04, A06, C02, and C08. Their increased methylation started from F_1_ and was then maintained from F_2_ to F_8_. Transcriptome analysis showed that genes located in both types of regions showed no significant expression changes.

We also identified 17 genomic regions (corresponding to one genomic locus in 17 samples) with decreased methylation. However, this contrasts with our common understanding of the relationship between DNA methylation and gene expression in the way that they also showed no effect on the expression of genes located in these regions. Moreover, in this genomic locus with decreased methylation, there was a much higher ratio of LTR-transposons, especially of Gypsy elements. The stable decrease in DNA methylation among progeny may indicate the anti-methylation property of this region. Potentially, the high abundance of LTR/Gypsy in this region may be activated under a low level of DNA methylation and start to boost and insert into other genomic regions after tetraploidization.

### Eventual Methylation Decrease Accompanies the Evolution of Polyploids

We observed a decrease in whole genome methylation after synthetic tetraploidization, and even after tens of thousands of generations as inferred from the decreased methylation levels in the neo-tetraploid (domesticated) *B. napus*. The consecutive decrease in methylation level was observed in the synthetic tetraploid genomes of *rapa* and *oleracea* from the second generation of self-crossing (F_4_)—a slight decrease in each generation compared to its parental generation. This methylation decrease trend was supported by the much lower methylation level in the natural neo-tetraploid genome of *B. napus*. As reported in previous studies (44–46), TE methylation suppresses the activity of frequently inserting into the host genome. However, the consecutive decrease in methylation may gradually relax the suppression effect on TE activity, which had adverse effects on the stability of the host genome. This subsequent TE activation released by the decreased methylation might then create an increasing number of genomic polymorphisms in the otherwise genetically fixed polyploid population and promote breakpoint-associated chromosomal duplications and other aberrations. This fanciful idea will be tested in future studies.

The long-time methylation decrease after polyploidization and its relationship with TE activity may also fit in to the “radiation lag” hypothesis (47), which refers to an argument that, following a polyploidy in the flowering plant phylogenetic tree, there is indeed a burst of diversity but only after a lag period. This hypothesis is not proved in any way. However, our data showing long-time methylation decrease fits the lag time before species radiation.

## MATERIALS AND METHODS

### Plant Materials and Sample Preparation

The *B. rapa* accession Chiifu (‘Chiifu-401-42’) and *B. oleracea* accession ‘JZS’ were selected as diploid parents to conduct the hybridization assay. After crossing the two parents, the hybrid immature embryos obtained from the excised siliques were rescued by culturing them on the MS medium to prevent the degradation caused by the reproductive barrier. The apical meristem of one survived hybrid F_1_ was cultured on MS medium with 0.02% colchicine for five days to induce the amphidiploid, which were further transferred onto the regeneration medium. One F_2_ with doubled chromosomes was successfully obtained. For the acquisition of F_3_–F_8_ generations, we collected seeds from the previous generation and planted them for the next round of self-pollination. The 6-8th fresh leaves were harvested from the eight-leaf stage seedlings of the diploid parents, hybrid F_1_, amphidiploid F_2_, F_3_–F_8_ progeny, and *B. napus* ZS. In particular, three individual plants were randomly selected from each generation of F_3_–F_8_ as biological replicates. All of the collected leaf samples were divided in half, flash frozen in liquid nitrogen, and separately stored at −80°C for RNA sequencing and bisulfite sequencing.

### Identification of Orthologous and Paralogous Genes

The genome versions of *B. rapa* Chiifu (v3.0) and *B. oleracea* JZS (v2.0), as well as corresponding gene annotations (v3.1 for BrChiifu and v2 for BoJZS, are available at http://brassicadb.cn/#/Download/) and were used as references to identify orthologous genes (38, 39). The syntenic orthologous genes between BrChiifu and BoJZS genomes were identified using SynOrths (48), which were considered the syntenic paralogous gene pairs (*rapa* vs. *oleracea*) in the newly synthesized allotetraploid. As for *B. napus*, the genome assembly and gene annotations of Zhongshuang 11 (ZS) were obtained from http://ocri-genomics.org/Brassia_napus_genome_ZS11/. Syntenic paralogous genes between the Bo and Br subgenomes were identified using SynOrths (48).

### RNA Extraction, cDNA Library Preparation, Sequencing, and Gene Expression Analysis

Total RNA was extracted from the collected leaf samples using the Dynabeads mRNA DIRECT Kit (Illumina, San Diego, CA, USA) and quantified and assessed for integrity using a NanoDrop spectrophotometer and an Agilent Bioanalyzer. The cDNA library was constructed with 50 μg RNA from each sample using the VAHTS mRNA-seq v2 Library Prep Kit (Illumina, San Diego, CA, USA). The libraries were sequenced to generate 150 bp paired-end RNA-seq reads by Biomarker Technologies Corporation (Beijing, Beijing, China). The sequenced reads were assessed by FastQC (available at https://qubeshub.org/resources/fastqc), and then low-quality and adapter bases/reads were trimmed or filtered out to obtain the high-quality clean reads using Trimmomatic (49) with default parameters. The clean reads from the parent plants (Chiifu and JZS) were aligned to the BrChiifu and BoJZS genomes, respectively, by Hisat2 (50), and the fragment counts of each annotated gene were calculated using FeatureCounts (51), while the clean reads derived from F_1_ to F_8_ were mapped to the merged reference genome (BrChiifu and BoJZS) to calculate the gene expression level. The ZS genome was used as a reference genome for the RNA-seq data of ZS. The final quantification results were used to calculate the TPM. To shield the effects of highly similar sequences of the *B. rapa* and *B. oleracea* genomes on gene expression, we extracted the fragments that were multiply aligned to the locations belonging to the orthologous gene pairs between BrChiifu and BoJZS, and the counts of these extracted fragments were further equally distributed between the corresponding gene pairs. For each syntenic gene pair, the number of cross-mapping fragments (*n*) was counted, which were then evenly assigned to two syntenic genes (0.5*n* for each gene). The additional fragment number was added to the fragment number of each gene resulting from the standard quantification process of FeatureCounts (51).

### Comparison of the Expression Levels of Gene Pairs

Genes were sorted according to their expression levels and evaluated by TPM. To determine the dominantly expressed one in a certain gene pair (orthologous/paralogous pairs), their TPM values were compared, and the relative expression fold change was calculated. Only gene pairs with the dominant expressed gene showing a TPM > 5 were retained for subsequent analysis.

### Whole-genome Bisulfite Sequencing and Data Analysis

Total genomic DNA was extracted from leaf samples using the New Plant DNA Kit (TIANGEN). After DNA concentration and integrity were detected, bisulfite sequencing libraries were constructed according to the manufacturer’s instructions using the EZ DNA Methylation-Gold^TM^ Kit (Zymo Research, Irvine, CA, USA). Bisulfite sequencing libraries were sequenced to generate 150 bp paired-end reads. The strategy for choosing the reference genome for different materials was the same as that described for RNA-seq analysis. Clean reads were generated by removing low-quality and adapter bases/reads using the same transcriptome filtering method. After the clean reads were aligned onto genomes, the Bismark tool (52) was used to remove duplicates and identify methylated loci. Considering that both CpG and CHG are symmetric methylations, the read counts (both methylated and unmethylated reads) of adjacent cytosine and guanine in the CpG and CHG loci were merged. The weighted methylation level of a certain region was calculated following a previously described method (53). To compare the methylation levels between the synthesized tetraploid genome and *B. napus* ZS, the methylation data of ZS were also aligned to the merged genome of BrChiifu and BoJZS, and similar analyses were conducted to determine the DNA methylation level.

### Resequencing and Data Analysis

Total genomic DNA was extracted from the same leaf samples used for bisulfite sequencing with the same method. The sequencing libraries were constructed using the TruSeq Nano DNA HT sample preparation kit for Illumina, and were sequenced on the Illumina NovaSeq platform. Clean reads were aligned to the selected reference genome using BWA mem (54) with default parameters. Samtools was used to sort the resultant BAM files and to call the read depth with the parameter ’-Q 5’.

### Profiling of Genomic Methylation Patterns

To visualize the methylation variations between different individual plants on a genomic scale, a 500 kb sliding window with a step set at 50 kb was used to scan the whole genome, and the weighted methylation level of each window was calculated. Genomic regions showing divergent methylation patterns were screened for changes of two-fold or greater.

### Determination of TE Composition and DNA Methylation Levels Surrounding TEs

The repeat sequences from *B. rapa* Chiifu, *B. oleracea* JZS, and *B. napus* ZS were predicted through both RepeatModeler (version 1.0.10) and RepeatMasker (version 4.0.3) (55). TE composition and content located at the increased and decreased methylation regions were extracted with in-house scripts. The DNA methylation level surrounding the TEs was calculated using each 100 bp sliding window with a step set at 10 bp. TE body was separated into 10 bins with an equal length and submitted to the window screening and methylation level calculation.

## ACKNOWLEDGEMENTS

This work was supported by the National Natural Science Foundation of China (NSFC grants 31972411, 31722048, and 31630068), Central Public-interest Scientific Institution Basal Research Fund (No.Y2022PT23), and the Science and Technology Innovation Program of the Chinese Academy of Agricultural Sciences, and the Key Laboratory of Biology and Genetic Improvement of Horticultural Crops, Ministry of Agriculture, P.R. China. MF supported by NIFA, US Department of Agriculture via UC-Berkeley, USA.

## Notes

### Competing Interest Statement

The authors have declared no competing interest.

