## Supplementary Figures for "Variations in Homoeologous Dosage and Epigenomics Mark the Early Evolution of Synthetic *Brassica* Tetraploids"

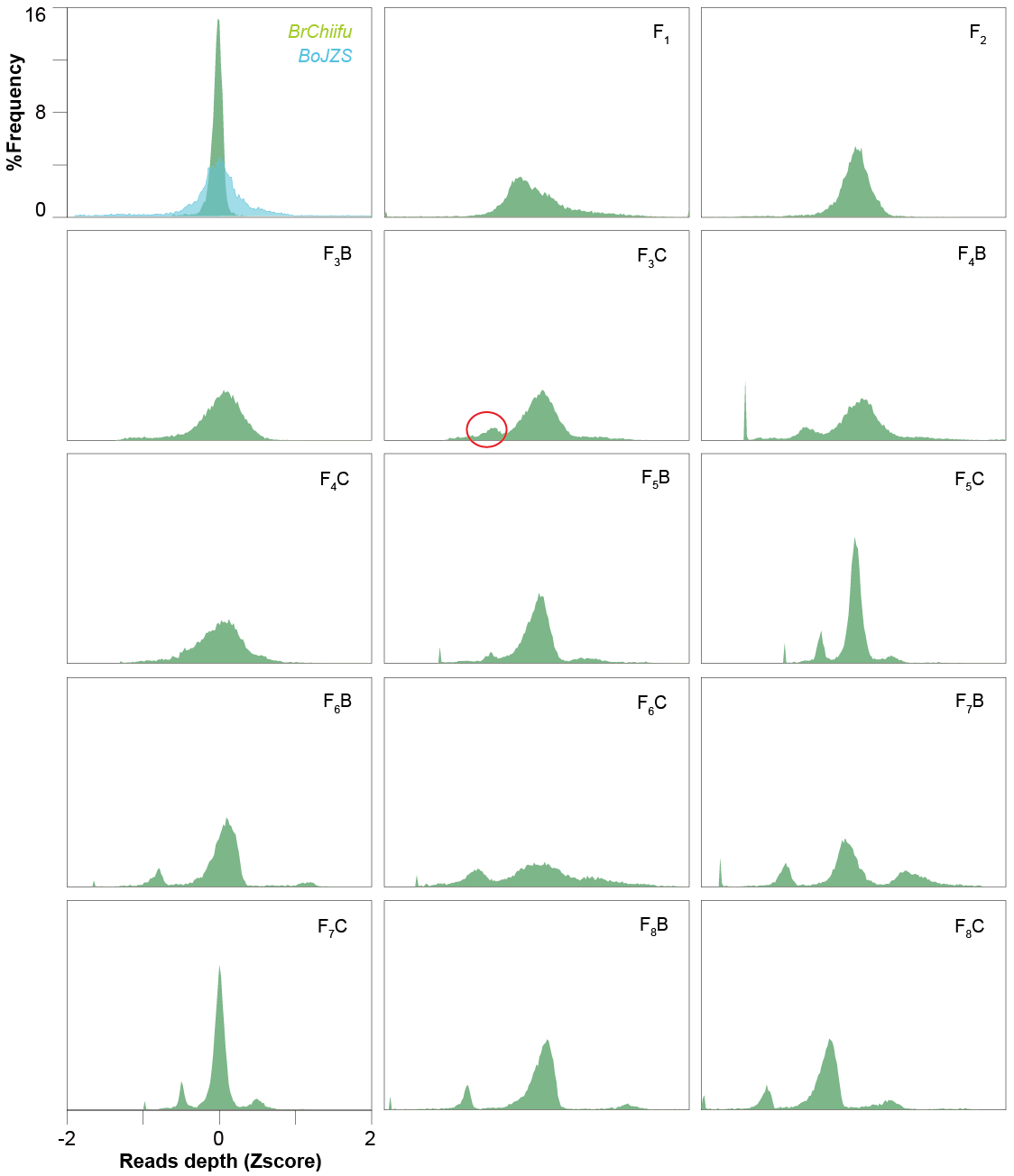

**Supplementary Figure 1.** The frequency distribution of reads depth in a 200 kb window sliding across the genomes of two parents and F_1_B/C to F_8_B/C.

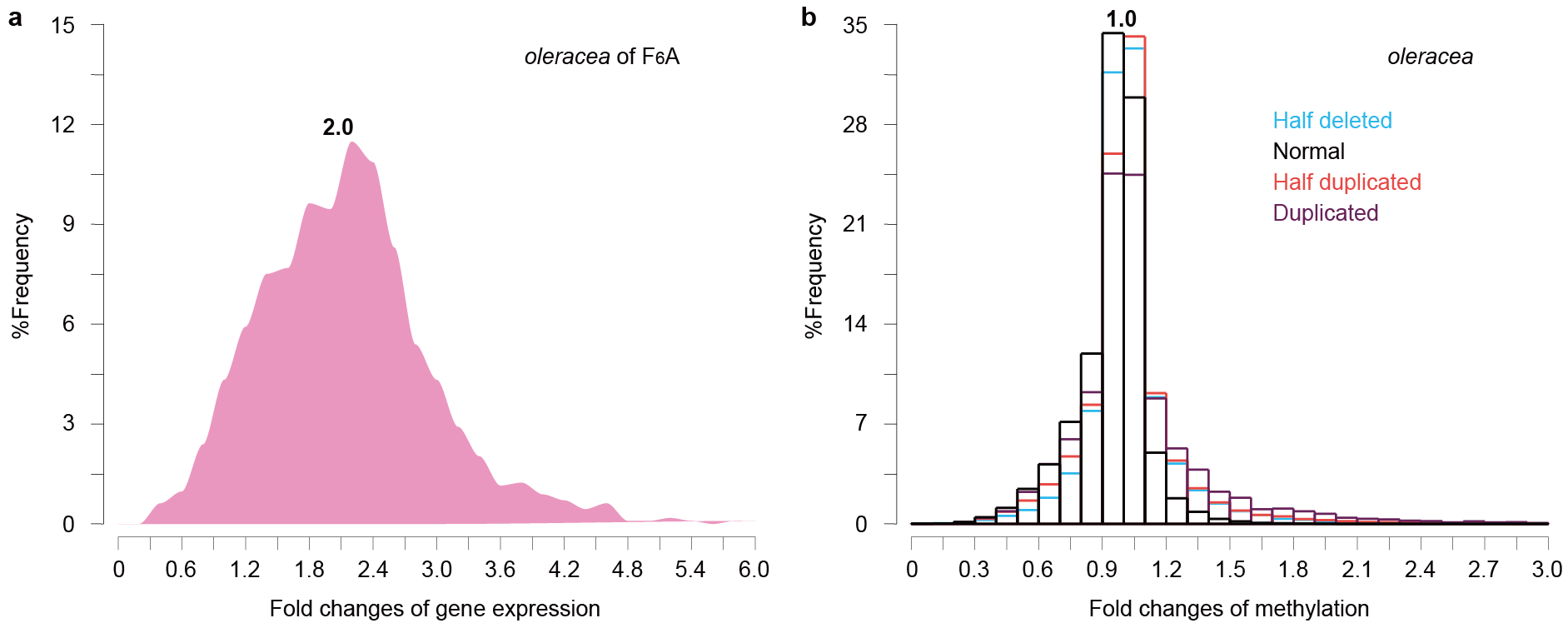

**Supplementary Figure 2. The frequency distribution of fold changes on expression and methylation of genes located at the large duplicated genomic regions.** (**a**) The frequency distribution of fold changes of gene expression in the duplicated genomic fragments. (**b**) The frequency distribution of fold changes of gene methylation in subgenome *oleracea*.

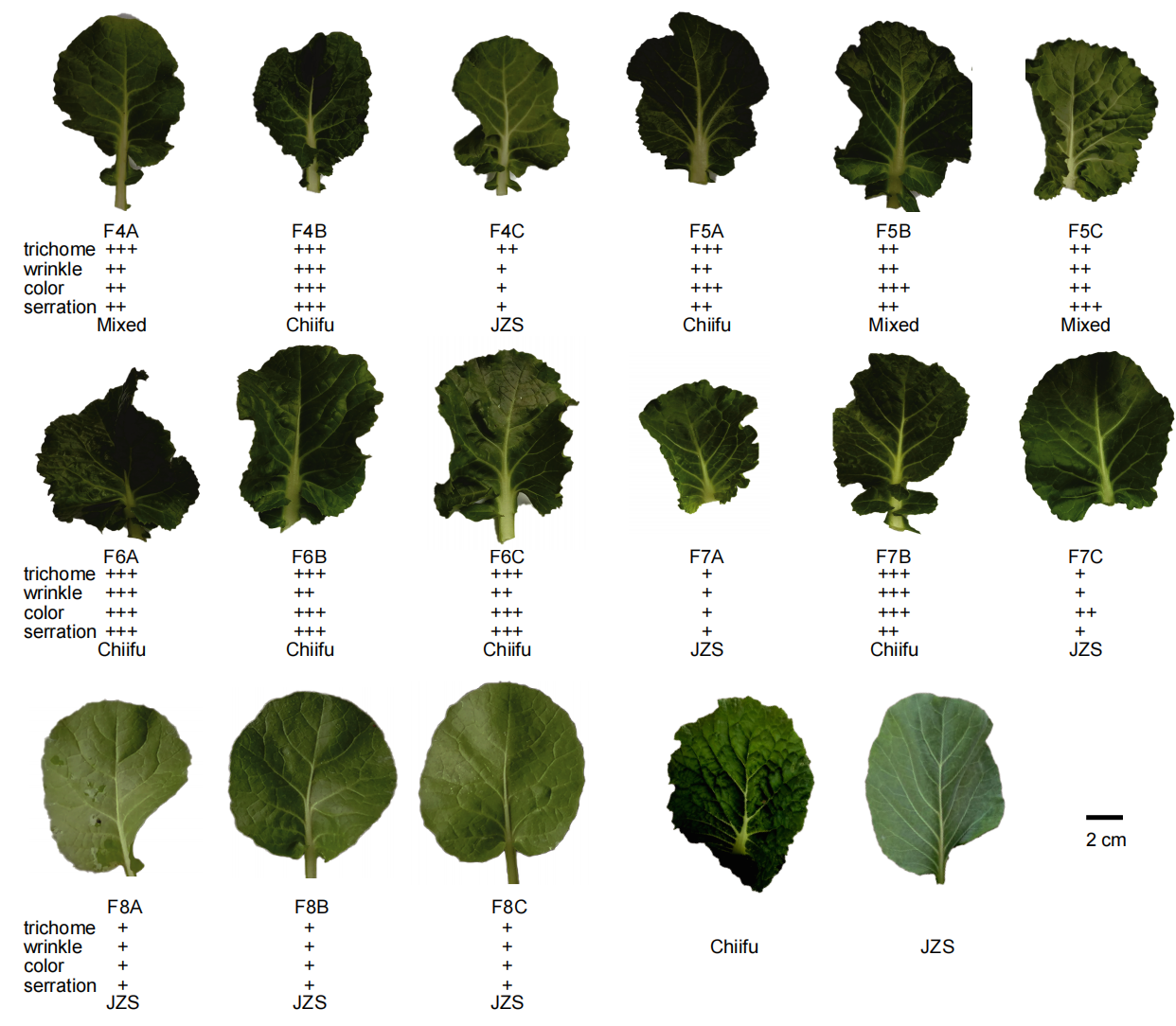

**Supplementary Figure 3. The leaf phenotype diversity of the three individual plants from F_4_ to F_8_ of the synthesized tetraploids.**

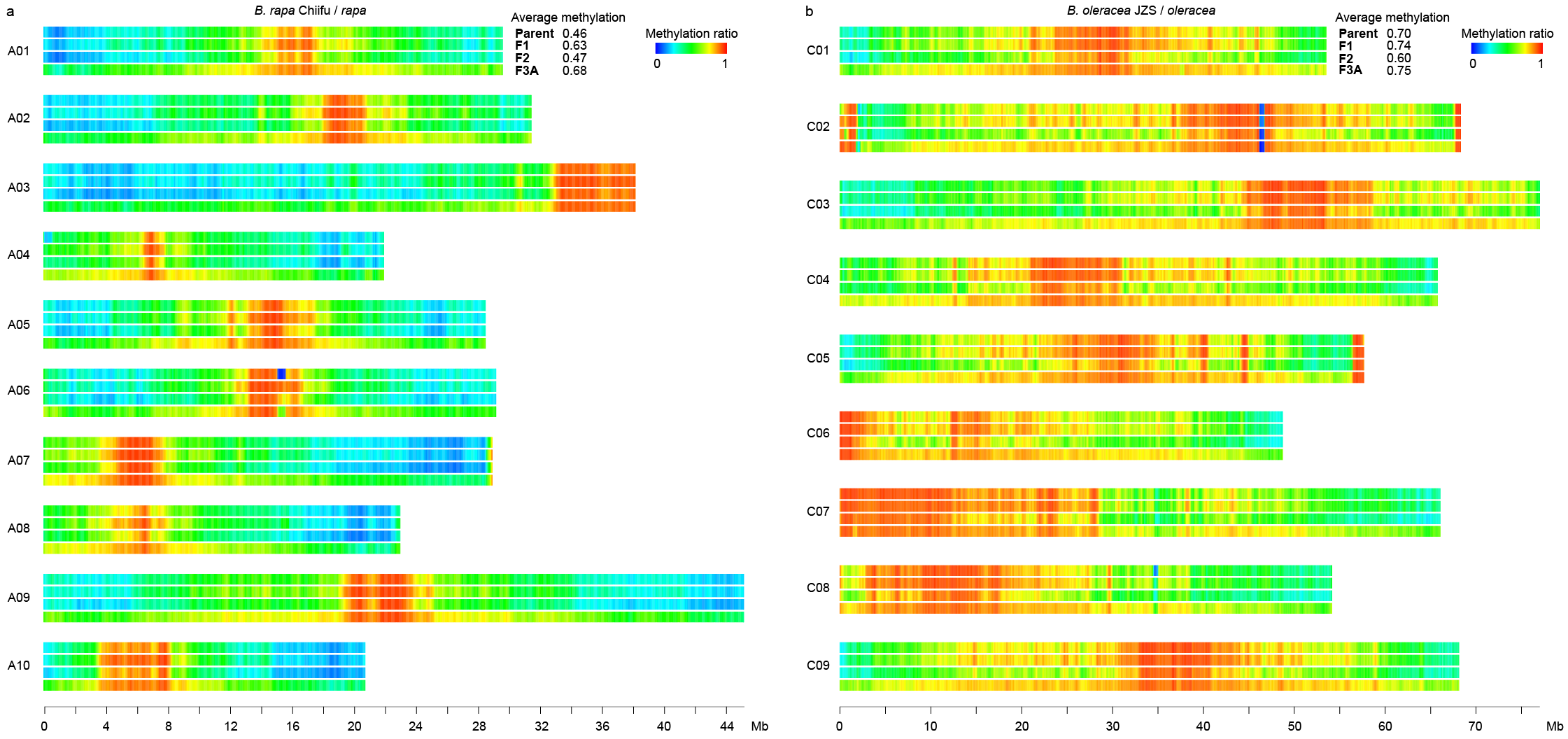

**Supplementary Figure 4. The heatmap of the methylation ratio of CpG loci across the parental genomes and the merged sub-genomes in F_1_, F_2_ and F_3_A.** (**a**) The heatmap showing the methylation ratio on genomes of the parent and the subgenome *rapa*; (**b**) The heatmap showing the methylation ratio on genomes of the parent and the subgenome *oleracea*.

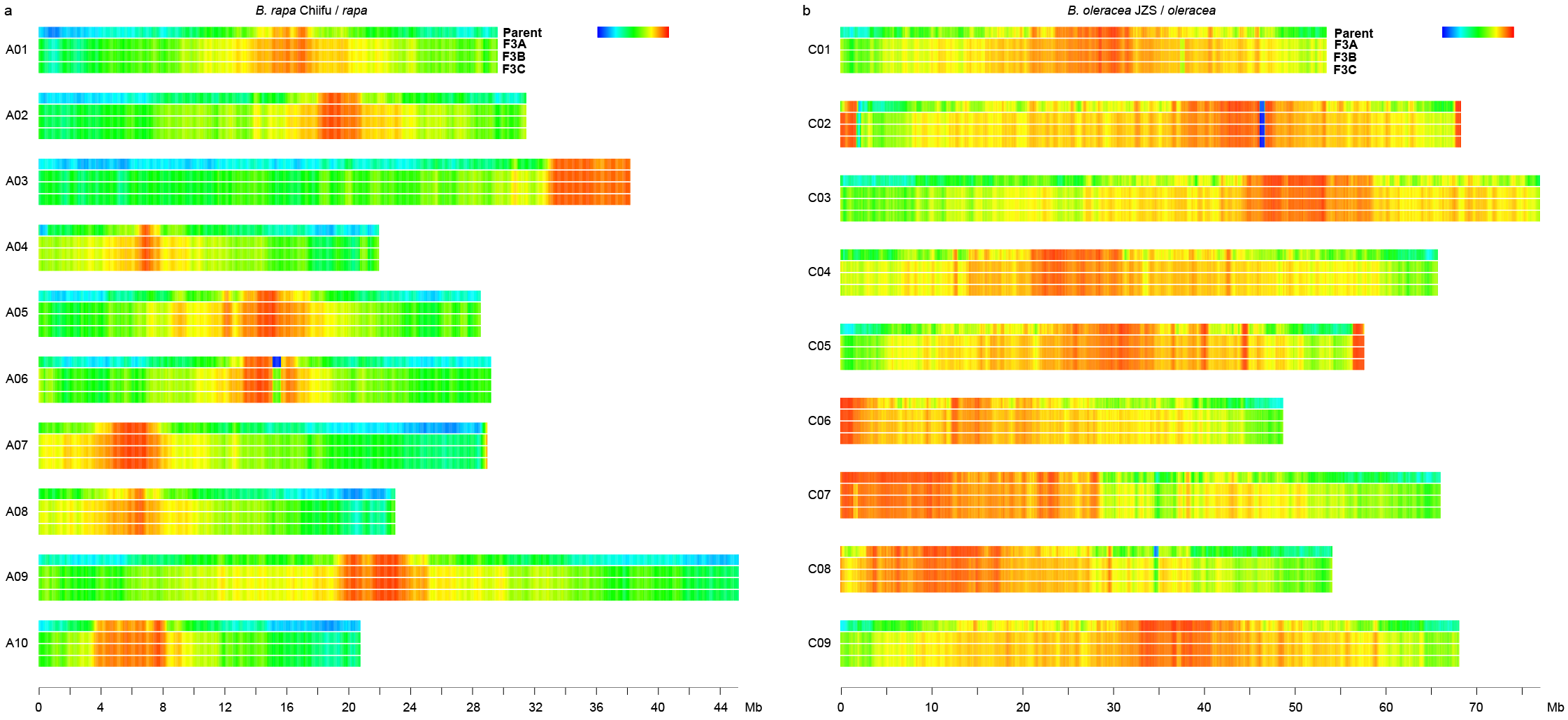

**Supplementary Figure 5. The heatmap of the methylation ratio of CpG loci across the merged sub-genomes in F_3_A, F_3_B, and F_3_C.** (**a**) The heatmap showing the methylation ratio on genomes of the parent and the subgenome *rapa*; (**b**) The heatmap showing the methylation ratio on genomes of the parent and the subgenome *oleracea*.

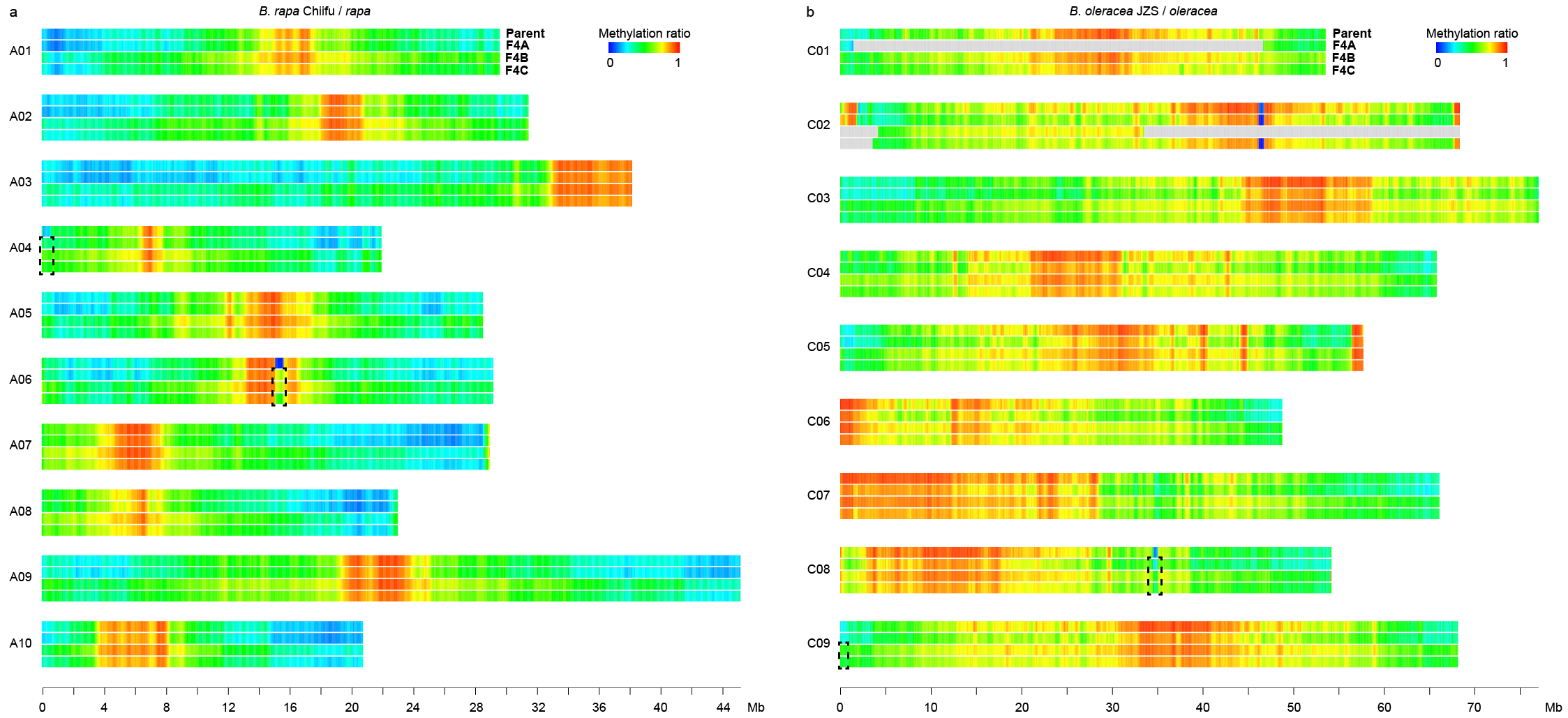

**Supplementary Figure 6. The heatmap of the methylation ratio of CpG loci across the merged sub-genomes in F_4_A, F_4_B, and F_4_C.** (**a**) The heatmap showing the methylation ratio on genomes of the parent and the subgenome *rapa*; (**b**) The heatmap showing the methylation ratio on genomes of the parent and the subgenome *oleracea*. The black dashed-line boxes mark the methylation increased or decreased regions; Grey bars denote genomic fragments that were deleted.

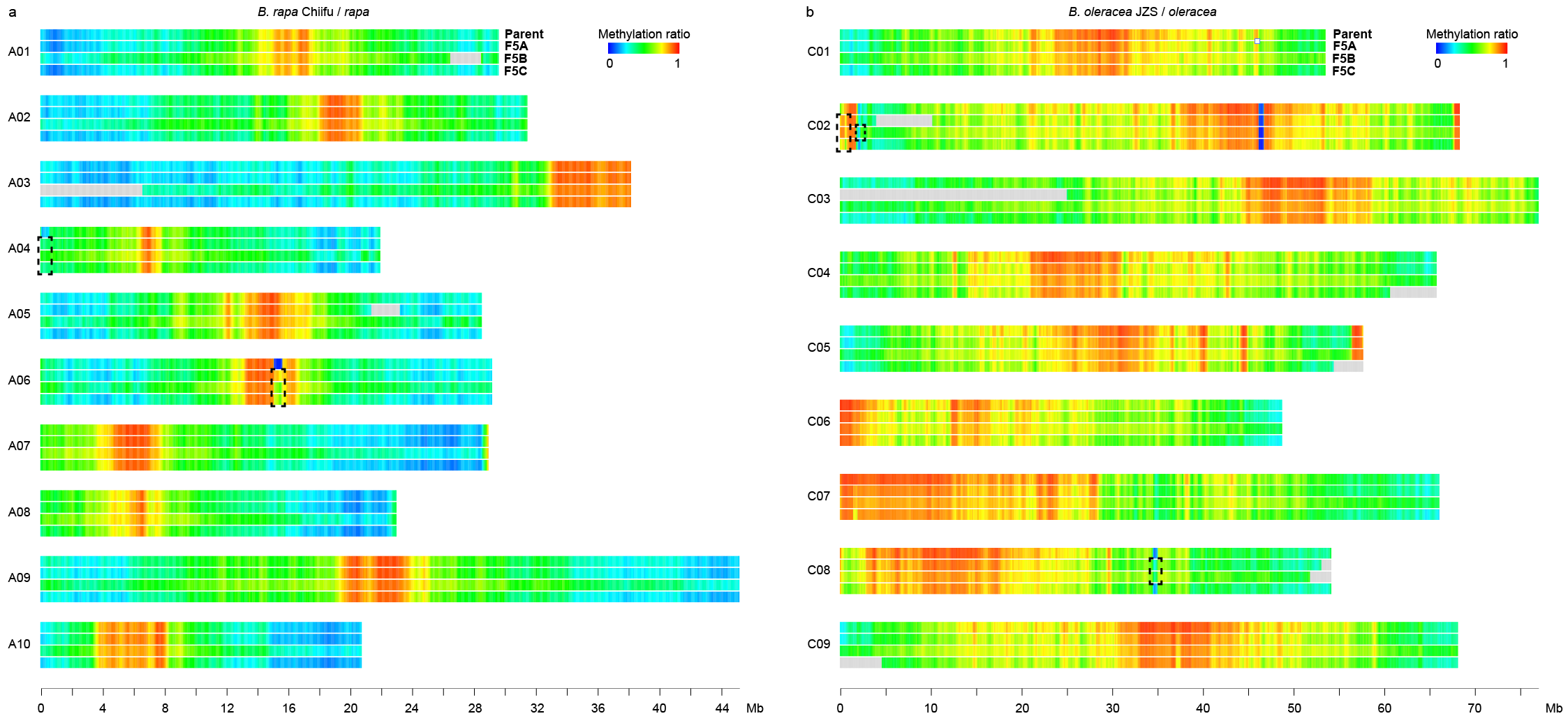

**Supplementary Figure 7. The heatmap of the methylation ratio of CpG loci across the merged sub-genomes in F_5_A, F_5_B, and F_5_C.** (**a**) The heatmap showing the methylation ratio on genomes of the parent and the subgenome *rapa*; (**b**) The heatmap showing the methylation ratio on genomes of the parent and the subgenome *oleracea*. The black dashed-line boxes mark the methylation increased or decreased regions; Grey bars denote genomic fragments that were deleted.

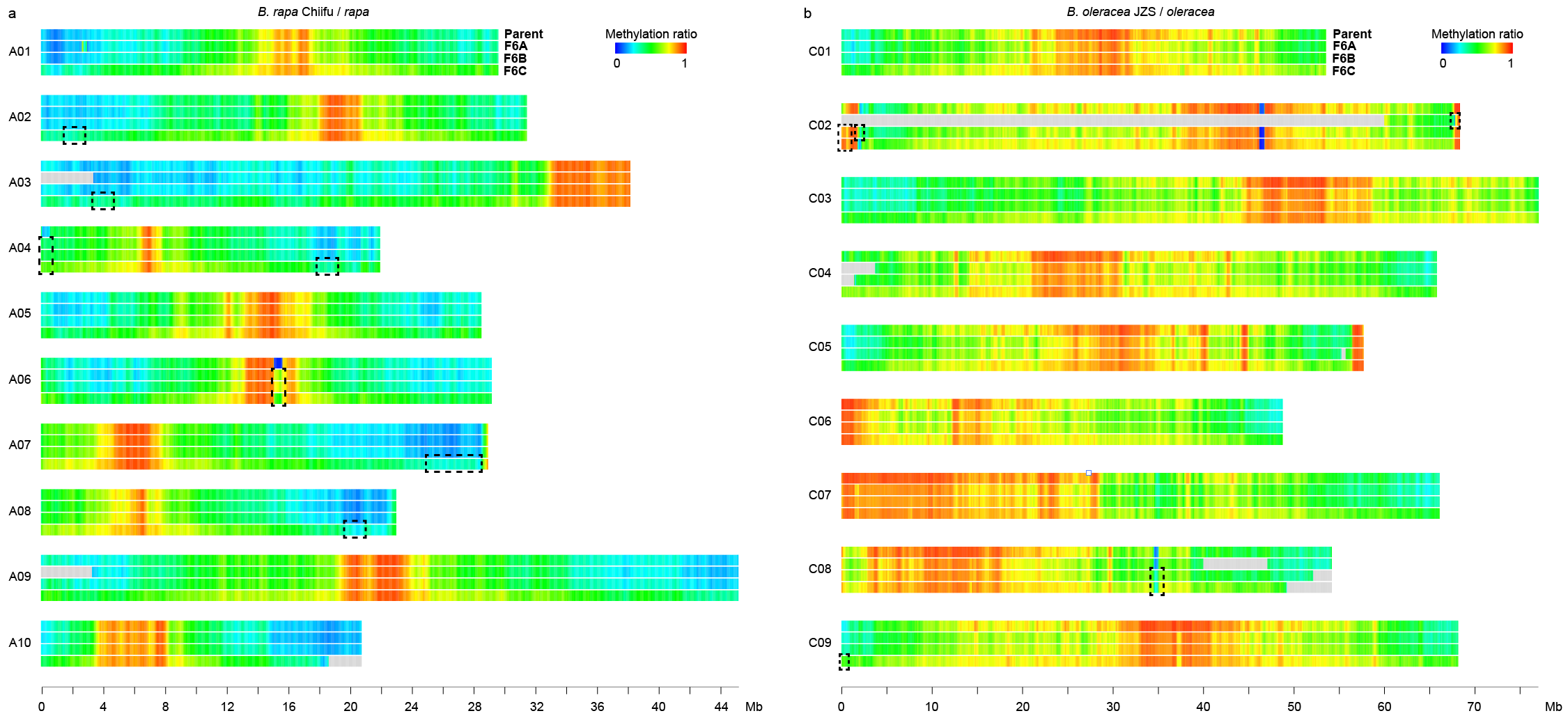

**Supplementary Figure 8. The heatmap of the methylation ratio of CpG loci across the merged sub-genomes in F_6_A, F_6_B, and F_6_C.** (**a**) The heatmap showing the methylation ratio on genomes of the parent and the subgenome *rapa*; (**b**) The heatmap showing the methylation ratio on genomes of the parent and the subgenome *oleracea*. The black dashed-line boxes mark the methylation increased or decreased regions; Grey bars denote genomic fragments that were deleted.

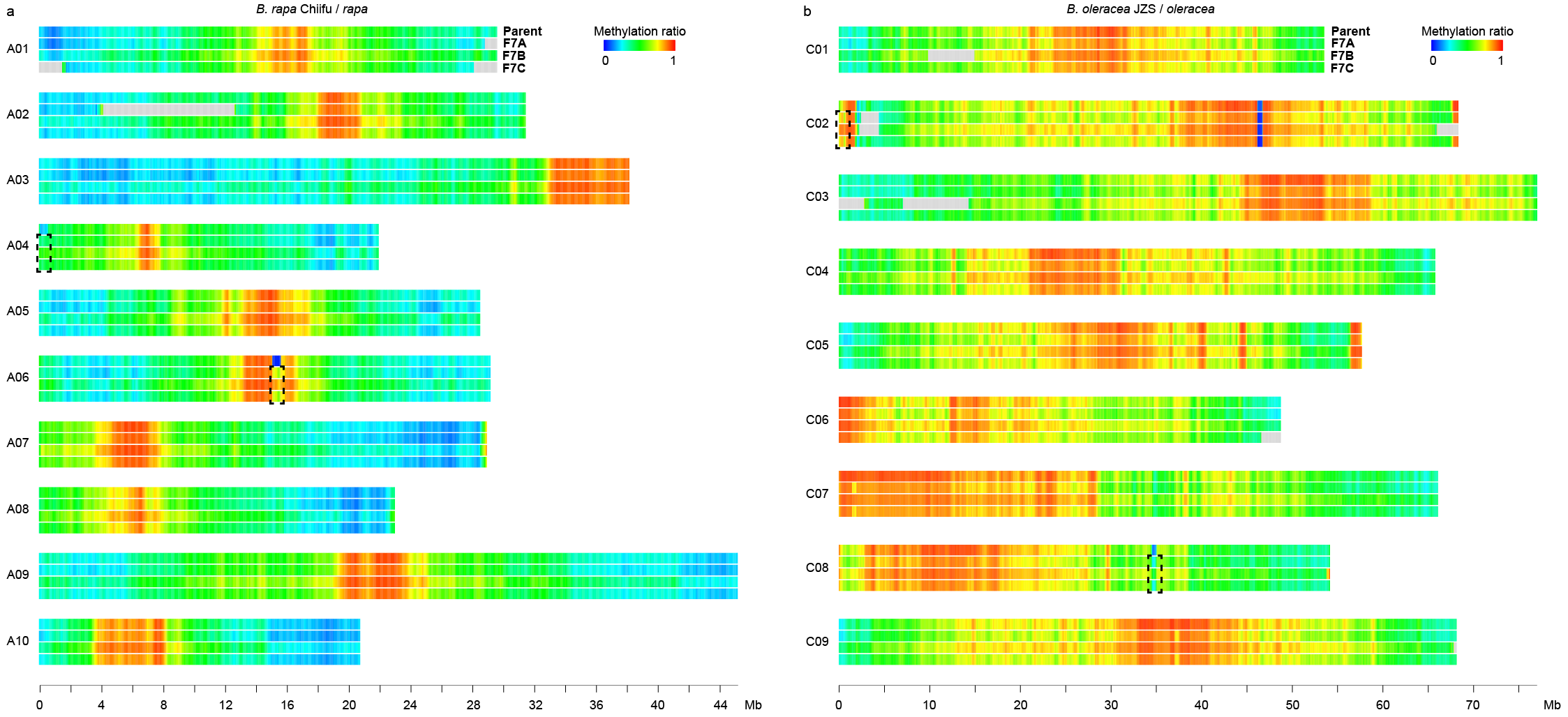

**Supplementary Figure 9. The heatmap of the methylation ratio of CpG loci across the merged sub-genomes in F_7_A, F_7_B, and F_7_C.** (**a**) The heatmap showing the methylation ratio on genomes of the parent and the subgenome *rapa*; (**b**) The heatmap showing the methylation ratio on genomes of the parent and the subgenome *oleracea*. The black dashed-line boxes mark the methylation increased or decreased regions; Grey bars denote genomic fragments that were deleted.

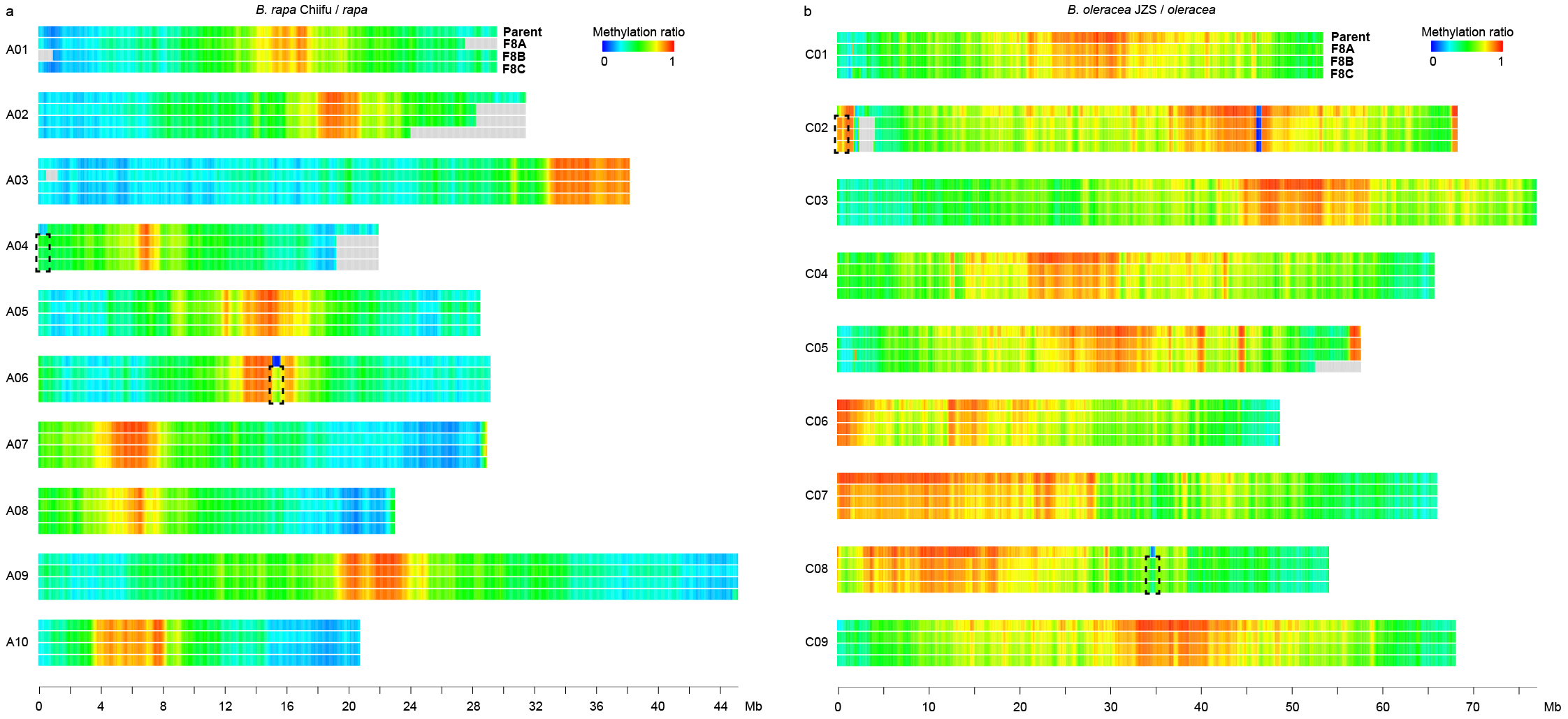

**Supplementary Figure 10. The heatmap of the methylation ratio of CpG loci across the merged sub-genomes in F_8_A, F_8_B, and F_8_C.** (**a**) The heatmap showing the methylation ratio on genomes of the parent and the subgenome *rapa*; (**b**) The heatmap showing the methylation ratio on genomes of the parent and the subgenome *oleracea*. The black dashed-line boxes mark the methylation increased or decreased regions; Grey bars denote genomic fragments that were deleted.
